# Genomic organization of *Trypanosoma cruzi* tRNA genes

**DOI:** 10.1101/2023.12.15.571829

**Authors:** Florencia Díaz-Viraqué, Ricardo Ehrlich, Carlos Robello

## Abstract

An often neglected class of genes in large-scale genome analyses is the non-protein-coding genes. In particular, due to the incompleteness of the genome assembly, it had not been possible to determine the genomic organization of the tRNA genes in *Trypanosoma cruzi* —a eukaryotic unicellular pathogen causative of disabling human Chagas disease—. Here, we analyzed the genic content and distribution of tRNA genes in the nuclear genome of different strains compared with the genome organization of other related trypanosomatids. We found synteny in most of the tDNAs clusters between *T. cruzi* and *Trypanosoma brucei*, demonstrating the importance of the genome location of these Pol III transcribed genes. The vast majority of the isoacceptor species are encoded by two genes, except for tDNA^SeC^, which is present as a tandem of 11 copies in the core compartment, associated with well-positioned nucleosomes. Finally, we describe a group of tRNA genes located at chromatin folding domain boundaries, possibly acting as chromatin insulators in *T. cruzi*.

## Introduction

Non-coding RNAs, or RNA genes, encode functional RNA products (Griffiths-Jones 2007) that participate in a variety of distinct biological processes, including protein synthesis, regulation of gene expression, RNA metabolism, as molecular scaffolds for the assembly of protein complexes, and epigenetic regulation, among others (Mattick et al., 2006). Within these genes, transfer RNA genes (tDNAs) are essential genes highly transcribed by RNA pol III that encode tRNAs —one of the most ancient molecules in living systems (Phizicky and Hopper, 2010)— that present conserved structures and sequence composition. tRNAs are between 73 - 93 nucleotide length and their secondary fold consists of four arms: DHU, TѱC, amino acid acceptor, and anticodon loops. Through the coaxial stacking of the DHU and TѱC loops, tRNA adopts an L-shaped tertiary structure. tRNA becomes loaded with an amino acid by its corresponding aminoacyl-tRNA synthetase enzyme. Then, the resulting aminoacyl-tRNA (aa-tRNA) plays a pivotal role as a substrate, actively participating in the peptide bond formation for protein synthesis. Discovered in 1958 (Hoagland et al., 1958), their only known function for decades was their role as decoders of the genetic code. Nowadays, it is known that in addition to their central role in translation, various very relevant extra-translational functions have been attributed to them in eukaryotes, prokaryotes, and archaea, such as regulation of transcription and translation, protein labeling for degradation, source of repetitive elements and small RNA fragments derived from functional tRNA molecules, chromatin domain barriers, DNA replication and cell death (Raina and Iba, 2014; Schimmel et al., 2018; Su et al., 2020; Ehrlich et al., 2021; Haldar and Kamakara, 2006).

In *Trypanosoma brucei* and *Leishmania major,* unicellular eukaryotes responsible for causing disabling human and animal diseases, most of the tRNA genes are organized into clusters of 2 to 10 genes separated by very short intergenic regions. These clusters contain other RNA polymerase III-transcribed genes and are confined in a subset of chromosomes (Tan et al., 2002; Padilla-Mejía et al., 2009). The genome organization of tRNA genes in *Trypanosoma cruzi*, the causative agent of Chagas disease, remains practically unknown until relatively recently due to its highly fragmented genome assembly. When the genome assemblies of *T. cruzi*, *T. brucei,* and *L. major* were published using short-read-based sequencing technologies (El-Sayed et al., 2005; Berriman et al., 2005; Ivens et al., 2005), the genome of *T. cruzi* remained highly fragmented and collapsed, mainly due to its intrinsic complexity: half of its genome is repetitive DNA (transposable elements, multigenic families, satellite DNA, and tandem repeats) (Pita et al., 2019); and several genes organized in tandem repeats. More recently, with the advent of long-read technology, several *T. cruzi* genomes assemblies have resulted using third-generation sequencing technologies (Berná et al., 2018; Callejas-Hernández et al., 2018; Díaz-Viraqué et al., 2019; Wang et al., 2021; Hoyos Sanchez et al., 2023). These approaches that improved the continuity and integrity of assemblies, coupled with extensive efforts in gene annotation, have expanded the number of coding genes and repetitive sequences annotated, providing a better estimation of gene expansion of the *T. cruzi* genome. Besides, it was described that the *T. cruzi* genome exhibits two different compartments (Berná et al., 2018), which harbors functional consequences at the level of three-dimensional chromatin organization (Díaz-Viraqué et al., 2023), allowing to explore the organization of genes with a functional perspective.

In this work, we study the genomic organization of tRNA genes in *T. cruzi*, analyze their genome context, provide a comprehensive comparative view of tRNA gene composition and organization with other trypanosomatids, and evaluate their implications in chromatin organization.

## Results

### ● Organization of tRNA genes in Trypanosoma cruzi

To examine the complete set of tRNA genes in the *T. cruzi* Dm28c genome, we searched these genes using tRNAscan-SE and studied their characteristics and genome context. *T. cruzi* contains 105 tRNA genes and four pseudogenes (**Figure 1** **and Supplementary Table 1)**; the complete representation of the genomic organization of tDNAs, including their flanking protein-coding genes, is shown in **Supplementary** Figure 1. Most are organized into clusters: 84 genes are organized in 20 clusters, and 21 are single genes (**Figure 1**). As in *L. major* and *T. brucei* (Padilla-Mejía et al., 2009), the number of tRNA genes per cluster ranges from 2 to 10, and none of the clusters is composed of isoacceptors. An intriguing feature is that half of the adjacent tDNA pairs in clusters have antiparallel orientation, while the majority in parallel orientation is the most common organization in other organisms (Bermudez-Santana et al., 2010). Although gene expansion through gene duplication is a common characteristic of the *T. cruzi* genome (Jackson, 2015), no tDNA duplication was found, except for the selenocysteine tRNA genes (described below).

**Figure 1:**
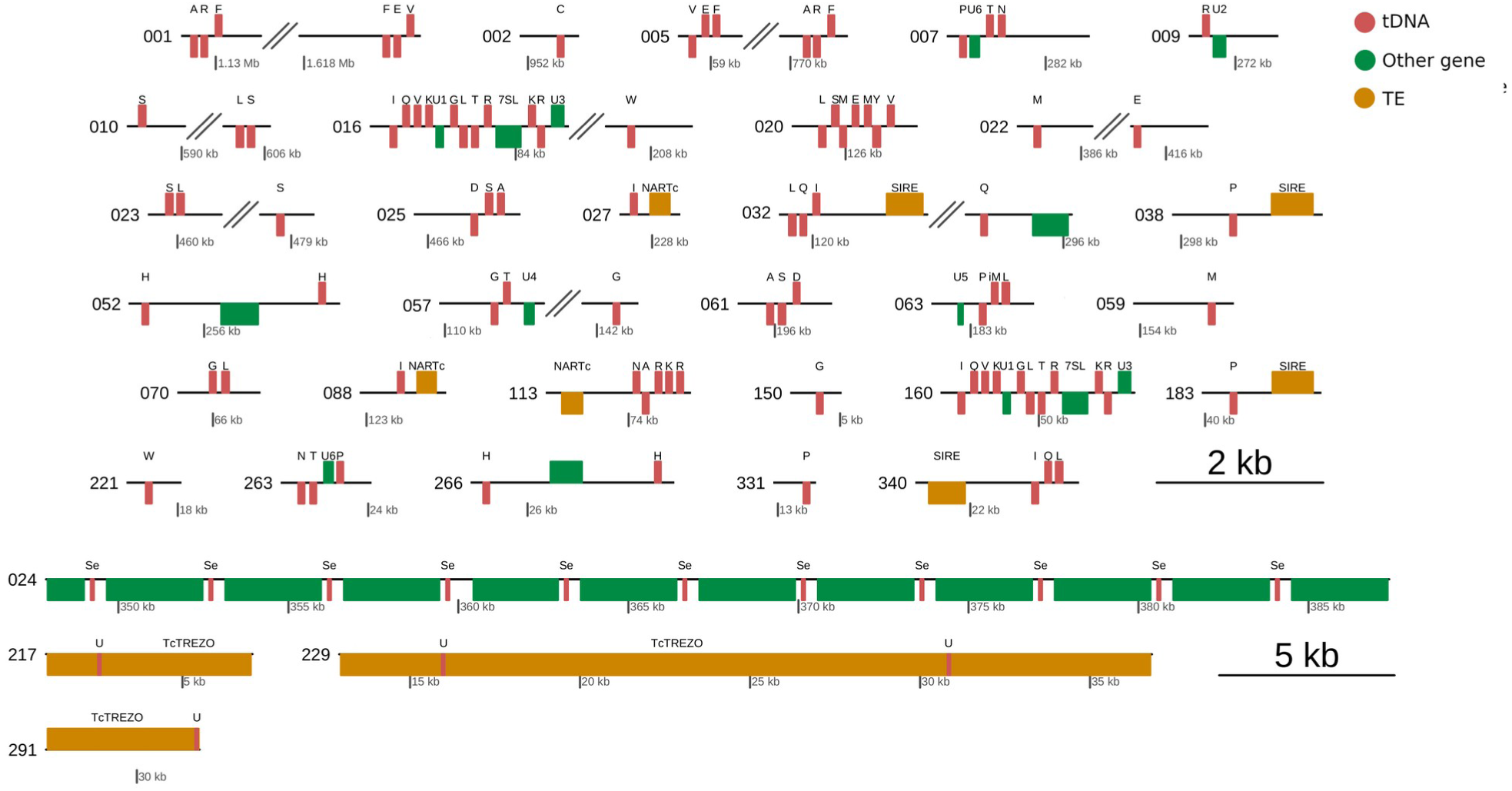
tDNA genomic loci in *T. cruzi*. tDNA coding genes annotated using tRNA-scan SE were identified in 40 different loci in the *T. cruzi* genome. 77% of the tDNAs are organized in clusters, and linear genome association with other Pol III transcribed genes is similar to *T. brucei*. tDNA loci are also composed of 7SL genes and snRNAs, while ribosomal genes are absent. Protein-coding genes are indicated in green without annotation. U: tRNA undetermined/unknown isotypes/pseudogenes; Se: Selenocysteine tRNAs.

The 109 tDNAs are located on 40 loci on 30 different scaffolds (51 tDNAs in core compartment, 7 in disruptive, and 51 in undetermined contigs), and 33% of the tDNA loci present transposable elements (TE). There are 14 TE within the tRNA gene loci, 9 are SIRE elements, and 5 are NARTc **(Supplementary** Figure 1). The four pseudogenes, with undetermined anticodon, are found in tandem with the transposable retroelement TcTREZO (**Figure 1**). Moreover, all tDNA genomic loci are surrounded by several TE fragments **(Supplementary** Figure 2) and repeat elements.

Strand-switch regions (SSRs) have been described as the genomic regions where several tRNA genes are located in *L. major* and *T. brucei* (Padilla-Mejía et al., 2009). In *T. cruzi*, we found that 52% of tRNA genes (excluding selenocysteine tRNA and pseudogenes for their particular organization) are located in these SSRs, either in clusters (9 clusters) or as single genes (10 tRNA genes). 11 of these 19 loci are present at divergent SRR —6 single genes and 15 genes are in 5 clusters— while the other 8 loci —4 single genes and 24 genes in 4 clusters— are found at convergent SSRs (**Supplementary** Figure 1).

As expected in eukaryotes, including trypanosomes (Kessler et al., 2018), tRNA^Tyr^ is an intron-containing tRNA. This explains why it is one of the largest genes, with those containing variable loop, tRNA^SeC^, tRNA^Ser^ y tRNA^Leu^ **(Supplementary Table 2 and Supplementary** Figure 3A and B). As in other kinetoplasts, tRNA^Tyr^ is the only intron-containing tRNA in the *T. cruzi* genome. However, this intron is two nucleotides longer and presents sequence divergence with the intron-tRNA^Tyr^ characterized in *Leishmania spp*. and *T. brucei* (Rubio et al., 2013).

### ● Expansion of selenocysteine tRNA genes in T. cruzi

Since selenocysteine tRNA (tRNA^SeC^) genes are not reported in *T. cruzi* genome annotations, three different algorithms were used to annotate them (see M&M). 11 tRNA^SeC^ genes were found in the Dm28c genome organized in tandem on a region of gene duplication, interspersed by a serine/threonine_protein_kinase coding gene (C4B63_24g205 to C4B63_24g214) (**Figures 1 and 2**). Given that RNA polymerase II transcribes these genes in *T. brucei* (Aeby et al., 2010), it seems reasonable that the same polymerase transcribes them in *T. cruzi,* which explains the linear genome association with other RNApol II transcribed genes. Due to the surprisingly high number of tRNA^SeC^ genes found, we searched this gene in other trypanosomatids genomes to study whether this gene expansion is a general mechanism in these organisms or exclusive of *T. cruzi*. We found one gene per *Leishmania* genome, but most Trypanosoma genomes present two or more copies **(Supplementary Table 3)**. Moreover, most mismatches in the multiple sequence alignment of these genes distinguish the sequences according to Leishmania or Trypanosoma genus (**Figure 2** **and Supplementary** Figure 4). Comparing the Trypanosoma and Leishmania sequences of tRNA^SeC^ with the sequences of other eukaryotes, we observed that the Trypanosoma sequences are the most distant **(Supplementary** Figure 5). These differences between genera do not seem to affect the secondary structure, but we cannot discard that this may determine different molecular interactions with proteins **(Supplementary** Figure 6). Besides presenting identical tRNA^SeC^ genes, similar protein-coding genes compose the locus in *T. cruzi* and *T. brucei* (**Figure 2**). These genes are interspersed by a “serine/threonine_protein_kinase” coding gene in *T. cruzi* and by “Calcium/calmodulin-dependent protein kinase, putative” in *T. brucei*. These kinases exhibit 44% identity at the amino acid level. Also, the first and last gene of the cluster is similar in both organisms: a methyltransferase and a hypothetical protein, which share 55% and 21% identity, respectively (**Figure 2**), whereas genes adjacent to tRNA^SeC^ in *L. major* are different.

**Figure 2:**
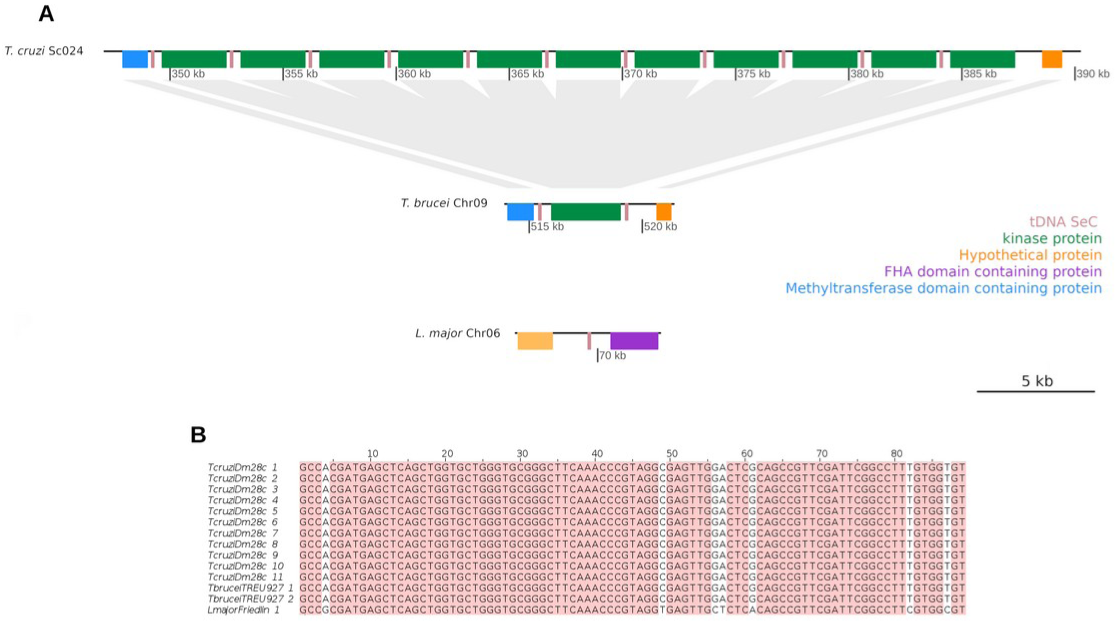
tRNA^SeC^ locus in *T. cruzi* (Dm28c), *T. brucei* (TREU927), and *L. major* (Friedlin) genomes. **A.** *T. cruzi* presents 11 copies of the tRNA^SeC^ gene while *T. brucei* has two copies and *L. major* only one. The tRNA^SeC^ genes in *T. cruzi* and *T. brucei* are interspersed by kinases, and this cluster is preceded by a hypothetical protein and followed by a protein containing a methyltransferase domain, whereas the genes on either side of tRNA^SeC^ in *L. major* differs. **B.** Multiple sequence analysis of tRNA^SeC^ genes. The genes of *T. cruzi* and *T. brucei* are identical, while the sequence of *L. major* presents seven differences from the previous ones.

### ● tRNA genes are syntenic in trypanosomes

Conservation of tRNA gene organization was previously considered scarce among trypanosomatids, and gene order was described as conserved in only a few cases (Padilla-Mejia et al., 2009). However, we found a high level of synteny in tRNA genomic loci among *T. cruzi* and *T. brucei* (**Figure 3A-F**). Of the 20 clusters of tRNAs, 19 (including tRNA^SeC^ genes) exhibit synteny with *T. brucei*. Another particularity we observed is that clusters are more dispersed in *T. cruzi* than in *T. brucei* genome. The conserved clusters are distributed in 16 *T. cruzi* scaffolds, while in *T. brucei,* tRNA genes are encoded in 6 chromosomes. In addition, although the identity and order of genes are exceptionally maintained, we found more frequently in *T. cruzi* some clusters interrupted by protein-coding genes or split in different scaffolds. These observations could be an artifact due to the highly fragmented genome assembly. In that case, this study can help to unite scaffolds, or this dispersion could indicate more chromosomal rearrangements in *T. cruzi*. There are three clusters interrupted by protein-coding genes exclusively in *T. cruzi*: the cluster encoded in scaffold PRFA01000032 (sc032) is interrupted by 175 Kb containing 52 protein-coding genes and 2 TE **(Figure 3A, second cluster)**; the cluster encoded in sc057 is interrupted by 33 Kb containing 21 genes (**Figure 3B, first cluster)**; and tRNA genes in Sc010 and Sc023 are separated by 16 and 19 Kb encoding 3 and 4 protein-coding genes, respectively, and 1 TE (**Figure 3D**). In the case of Chr11 and Sc005 (**Figure 3F**), there are two clusters of tDNAs with synteny encoded in two different loci. The linear distance between the clusters varies according to the species, being much longer in *T. cruzi* (711 Kb in *T. cruzi* vs. 17 Kb in *T. brucei*). In general, chromosomes in *T. brucei* contain more than one cluster or tRNA single genes. In contrast, in *T. cruzi,* most scaffolds code for only one locus of tDNA. Even clusters spaced in *T. brucei* are encoded in different scaffolds in *T. cruzi* (**Figure 3E**). In this case, the linear distance between these two clusters in *T. brucei* is 3172 bp, while one cluster is encoded in Sc061 and the other in Sc025 in *T. cruzi*. The relative positions of the tDNAs on these scaffolds, not in the extremes, imply that even though the assembly is very fragmented, the minimum lineal distance in the *T. cruzi* chromosome is much longer than in *T. brucei*. The same situation occurs in Chr10 and scaffolds Sc057 and Sc063 (**Figure 3B, first and second clusters)**, where the distance between the clusters is 40 kb in *T. brucei* while the minimum possible distance between the clusters in *T. cruzi,* taking the shortest distance to the ends, is 141 kb in *T. cruzi*.

**Figure 3:**
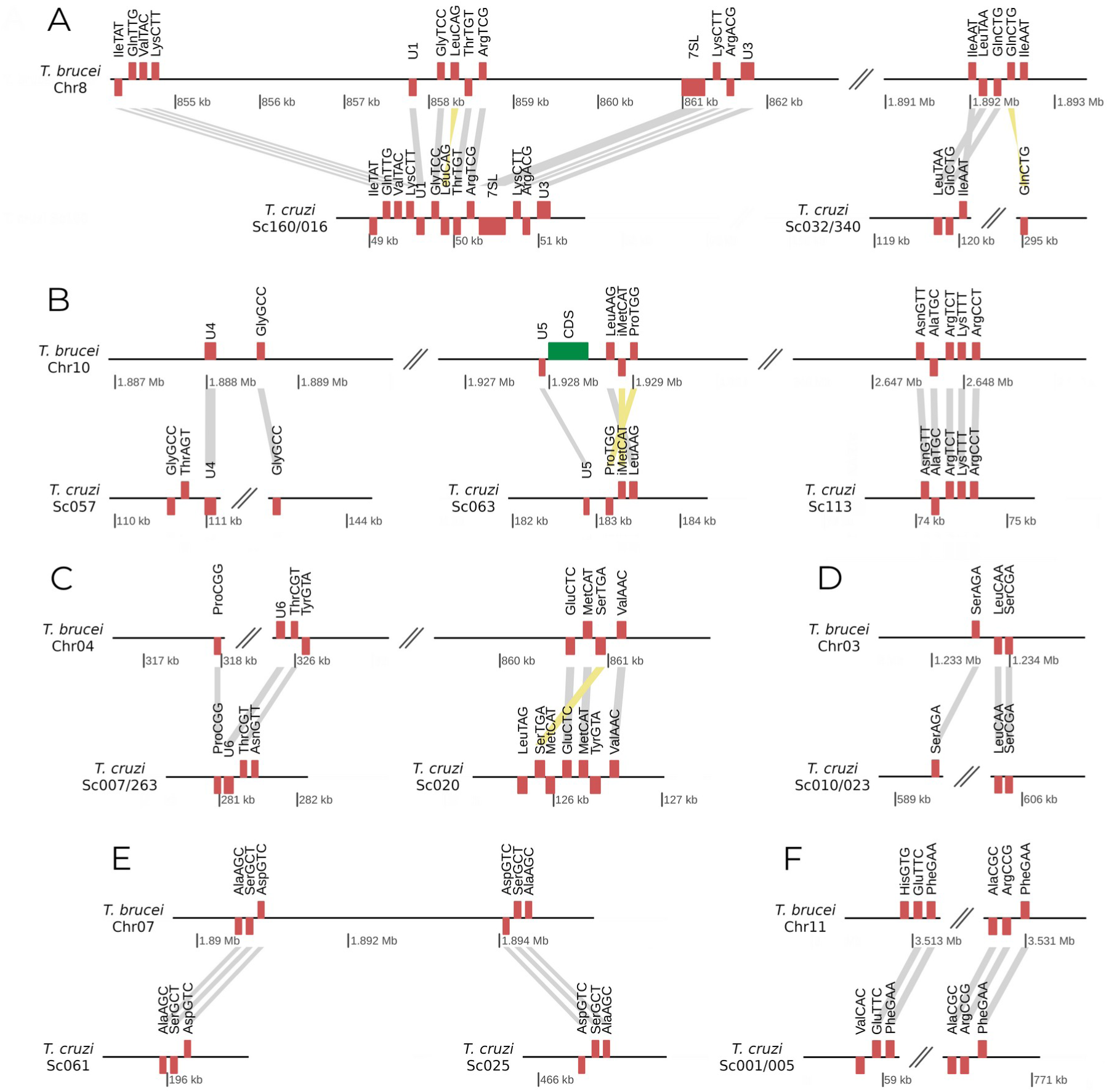
Syntenically conserved tDNAs between *T. cruzi* and *T. brucei*. Grey lines connect the orthologous genes in *T. brucei* chromosomes and *T. cruzi* scaffolds. In yellow are linked genes with an inverted relative position among the genes composing the tDNA locus.

Only the genomic organization of four clusters of *T. cruzi* had been previously reported, concomitantly to the synteny with *T. brucei* (Padilla-Mejia et al., 2009). The largest cluster (Sc016/160) was described as identical in gene composition but with an opposite orientation of three genes, tRNA^Leu^, 7SL RNA, and snRNA U1. As described below, we re-annotated these RNA genes and found that the sequence orientation of only one gene is distinct in this cluster. 7SL RNA and snRNA U1 were annotated in the wrong strand, tRNA^Leu^ being the only gene encoded in the opposite strand **(Figure 3A, first cluster)**. Another previously reported cluster is Sc032/Sc340 **(Figure 3A, second cluster)**. In this case, there is a difference in gene composition of the cluster between *T. cruzi* CL Brener strain used by Padilla-Mejia and collaborators (2018) and the *T. cruzi* Dm28c strain we analyzed. The cluster in Dm28c is Leu-Gln-Ile-Gln, while in CL Brener is Ile-Gln-Leu, due to the higher CL Brener genome fragmentation. Something similar occurs in scaffold Sc113 **(Figure 3B, third cluster)**, where the cluster in Dm28c is Asn-Ala-Arg-Lys-Arg (completely syntenic with *T. brucei*) while Ala gene is missing in CL Brener (Asn-Arg-Lys-Arg). To better examine this, we re-annotated tRNA genes in another hybrid strain: TCC **(Supplementary** Figures 7 and 8**, and Table 4**). The Ala gene is also present in TCC, as well as the association of locus containing LQI and Q.

### ● Isoacceptors, isodecoders, and codon usage

It was previously described that *T. cruzi* does not present a correlation between codon usage and the number of tRNA isoacceptors (Padilla-Mejia et al., 2009). To explore this, we compared these variables using all the 17197 annotated protein-coding genes in the non-hybrid strain Dm28c **(Supplementary Table 5)**. We observed no correlation between tRNA isoacceptor copy number and codon usage (Spearman correlation ρ = 0.19) (**Figure 4, upper panels)**. Therefore, the codon usage is not co-adapted with the relative abundance of tRNAs isoacceptors in *T. cruzi,* and this can not be explained by aneuploid. The genome of *T. cruzi* is partitioned into two large regions that correspond to chromatin compartments: the core (highly conserved and syntenic among trypanosomatids) and the species-specific disruptive compartment (synteny disruption), containing multigene families encoding for surface proteins (Berná et al., 2018). By eliminating genes from the disruptive compartment, the correlation between codon usage and the isoaccepting tRNA content increases (Spearman correlation ρ = 0.42) **(Figure 4, down panels)**, indicating different codon usage for the genes contained in each of the genome compartments. Besides, the correlation between the codon usage and tRNA isotypes on this non-hybrid strain remains invariant by removing genes from the disruptive compartment (Spearman correlation ρ = 0.73 and ρ = 0.72).

**Figure 4:**
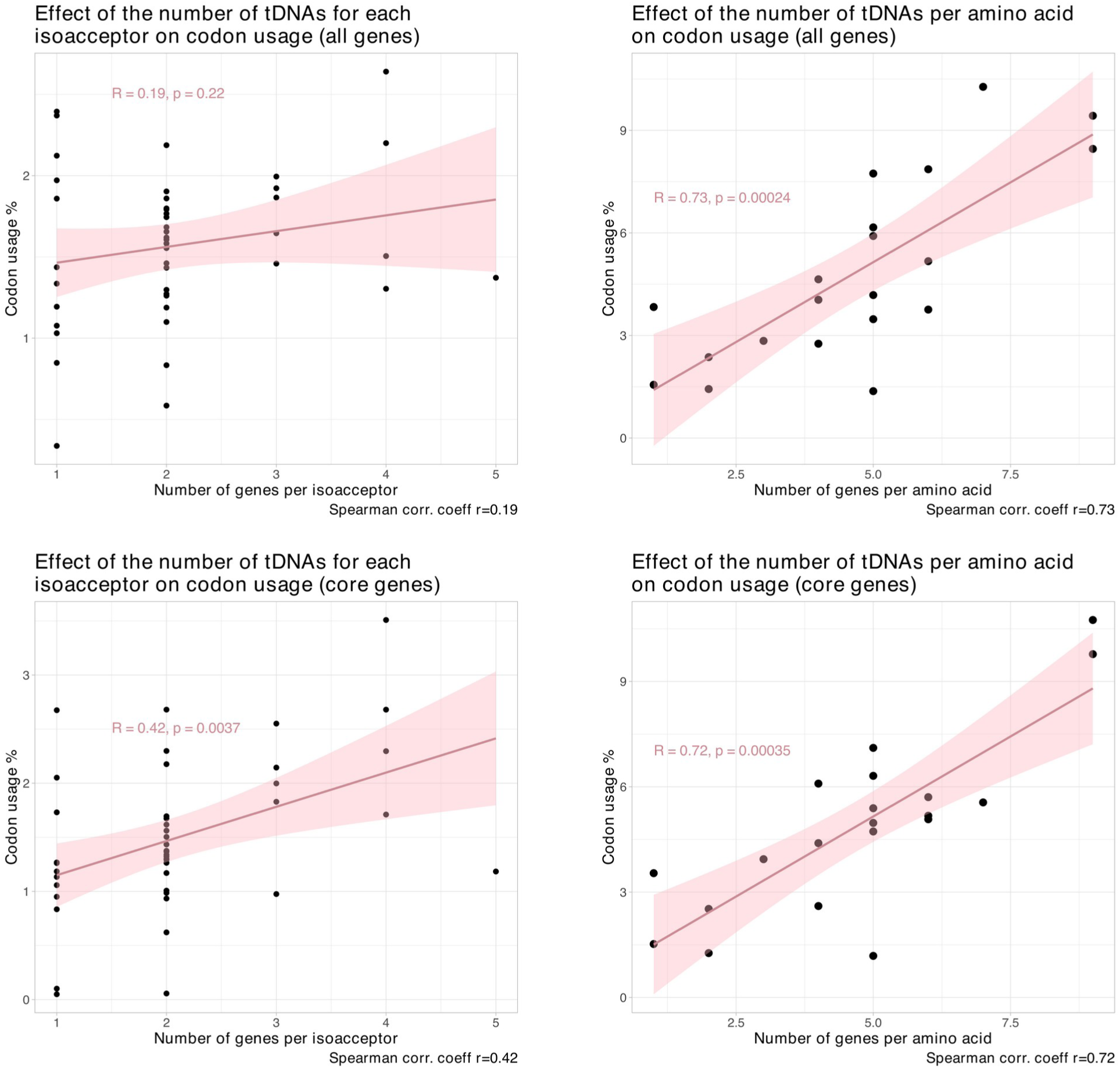
Correlation of codon usage with the number of tDNAs per amino acid and isoacceptor. Left panels: relation between codon usage and the number of tRNA genes for each isoacceptor. Spearman’s correlation coefficient is ρ=0.185516 and *p* value=0.2224. Right panels: Relation between codon usage and the number of tRNA genes per amino acid. Spearman’s correlation coefficient is ρ=0.7323246, and *p* value=0.0002413. The codon usage was calculated using the 17197 protein-coding genes annotated in the non-hybrid strain Dm28c (upper panels) and excluding the genes from the disruptive compartment (lower panels). Spearman’s correlation coefficient was used to measure the relationship between the variables.

### ● tRNA genes as chromatin insulators

tRNA genes are enriched at topologically associating domain (self-interacting genomic region) boundaries of the chromatin (Dixon et al., 2012), and in *T. cruzi* open chromatin was described to be developmentally regulated at tRNA gene loci (Lima et al., 2022). To further determine the relevance of *T. cruzi* tDNAs in chromatin organization, we annotated the tRNA genes in *T. cruzi* Brazil A4 strain, where chromatin folding domains (CFDs) were determined (Díaz-Viraqué et al., 2023). The 82 tRNA genes identified are located on 38 loci distributed on 15 chromosomes and 5 contigs (**Supplementary** Figure 9 and **Supplementary Table 6**). As in Dm28c, most of the tDNAs are organized into clusters: 55 genes (71%) are organized in 15 clusters, and 22 are single genes (**Supplementary** Figure 10). Interestingly, we found a practically identical genomic distribution in the tDNA clusters in Dm28c and Brazil A4 (**Figure 5**), which could be helpful for the improvement of the assembly of the genomes, as we analyze below.

**Figure 5:**
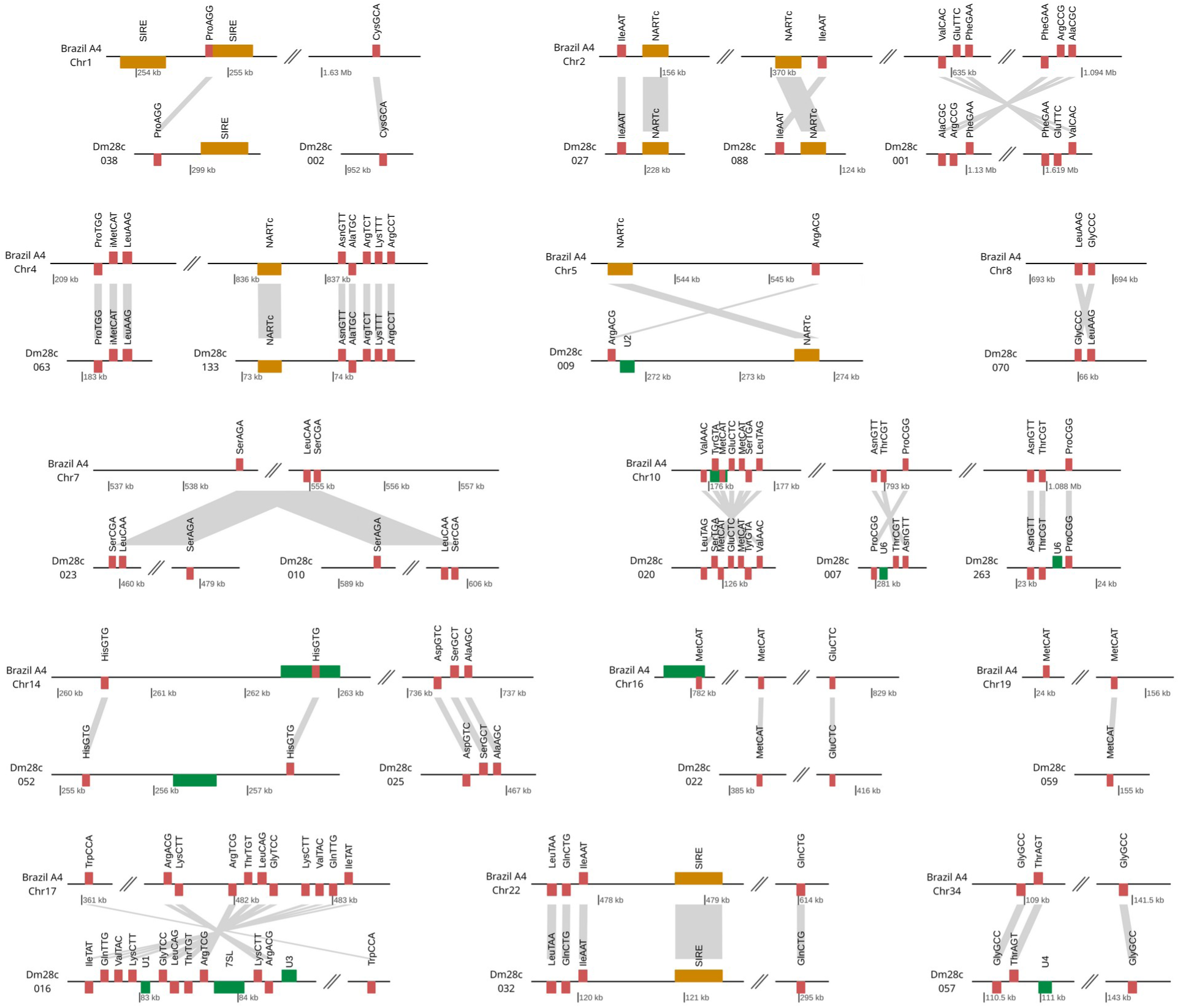
Syntenically conserved tDNAs between *T. cruzi* Dm28c and Brazil A4 strains. Grey lines connect the orthologous genes in Brazil A4 chromosomes and Dm28c scaffolds.

Regarding the genome organization, we found that 62 genes are present in the core compartment, 13 within the disruptive compartment, and seven are present in undetermined contigs enriched in transposable elements. We analyzed the chromatin domain boundaries, finding that 28 tDNAs are located at chromatin folding domains (CFDs) boundaries (**Figure 6A** and **Supplementary Table 7**). A particular case is the tRNA^SeC^ cluster, which, in addition to being close to a CFD boundary, there is a well-positioned nucleosome in each tRNA^SeC^ gene (**Figure 6B** **and Supplementary Table 8)**. Finally, we analyzed the protein-coding genes at the CFD boundaries, and of the 140 detected genes, we found an enrichment in surface genes and transposable elements **(Supplementary Table 9)**.

**Figure 6:**
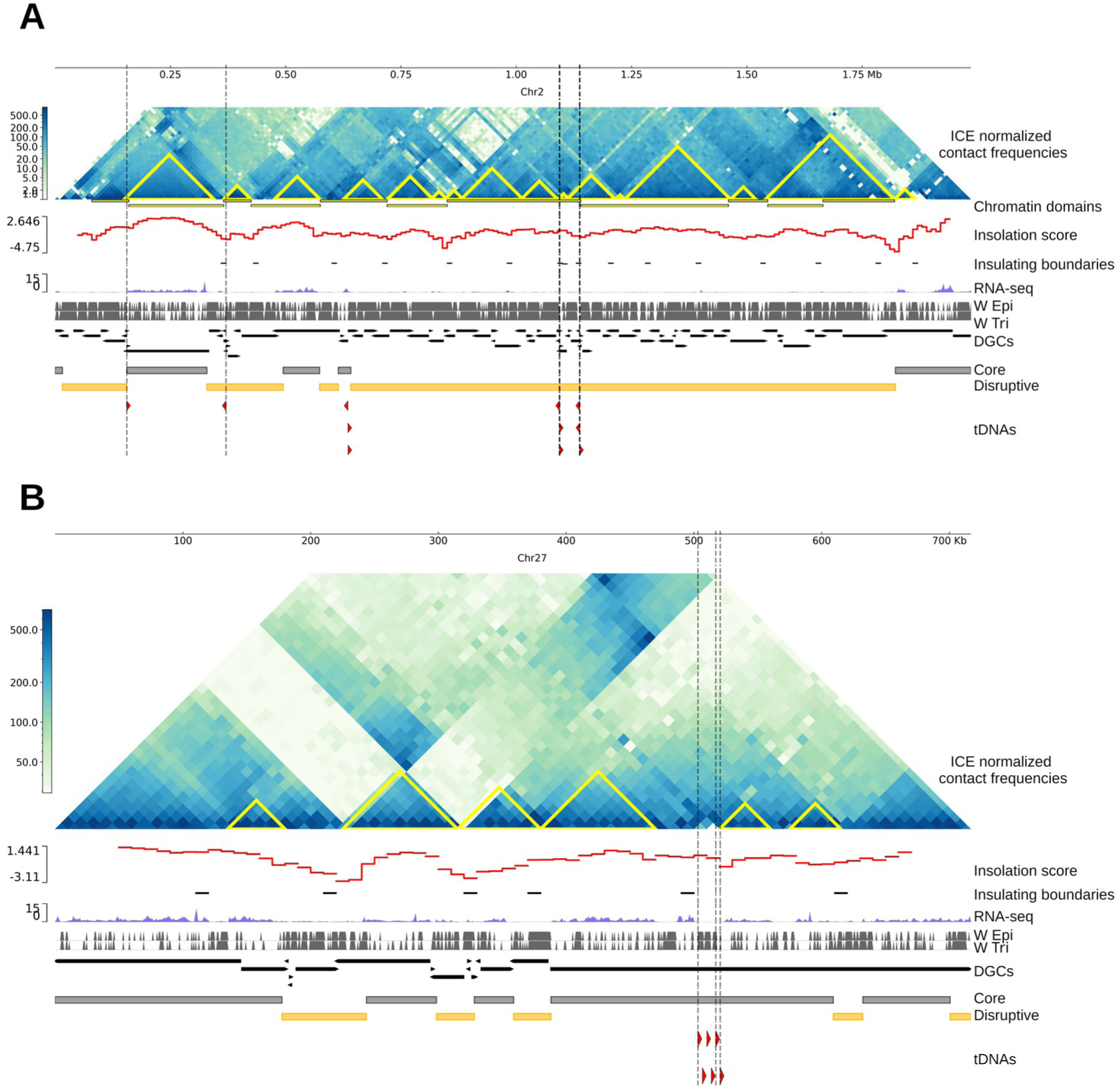
tRNA genes as chromatin insulators. Normalized (iterative correction and eigenvector decomposition, ICE) Hi-C interaction frequencies of chromosomes of *T. cruzi* Brazil A4 displayed as a two-dimensional heatmap at 10 kb resolution. Chromatin Folding Domains (CFDs) identified using HiCExplorer and TADtools (triangles on the Hi-C map) are indicated in yellow. Insulation score values were calculated using FAN-C, and the black horizontal dashes represent the local minima of the insulation score, indicatives of the region between two self-interacting domains. RNA expression across the chromosome is depicted as coverage plots using a bin size of 10 bp. Vertical grey bars indicate the position of all well-defined nucleosomes. Directional gene clusters (DGCs). The genomic position of core and disruptive genome compartments are represented in grey and yellow, respectively. tRNA genes identified as chromatin insulators are indicated with vertical dash lines. **A.** Normalized Hi-C interaction frequencies of chromosome 2. tRNA genes: tRNA^IleAAT^, tRNA^IleAAT^, tRNA^ValCAC^, tRNA^GluTTC^, tRNA^PheGAA^, tRNA^PheGAA^, tRNA^ArgCCG^, tRNA^AlaCGC^, tRNA^AlaCGC^, tRNA^ArgCCG^, tRNA^PheGAA^. **B.** Normalized Hi-C interaction frequencies of chromosome 27. tRNA genes: tRNA^SeCTCA^, tRNA^SeCTCA^, tRNA^SeCTCA^, tRNA^UndetNNN^, tRNA^SeCTCA^, tRNA^SeCTCA^.

### ● tRNA genes as markers for genome scaffolding

Genetic markers are regions of DNA that reside on the same chromosome and are co-inherited when not separated by recombination. These markers are used as reference points to identify and map different parts of a genome; therefore, they help improve genome assemblies, assign contigs to linkage groups (often chromosomes), and determine the correct order of contigs along chromosomes (Rice et al., 2019). To analyze whether tDNAs can be used as markers for genome scaffolding, we analyzed the synteny of the chromosomes of Brazil A4 and the scaffolds of Dm28c containing the tDNA clusters. Chromosome 1 from *T. cruzi* Brazil A4 presents synteny with two scaffolds of *T. cruzi* Dm28c genome assembly: 038 and 002 (**Figure 7**). Chromosome 7 from *T. cruzi* Brazil A4 presents synteny with two scaffolds of *T. cruzi* Dm28c genome assembly: 023 and 010. Chromosome 10 from *T. cruzi* Brazil A4 presents synteny with three scaffolds of *T. cruzi* Dm28c genome assembly: 020, 007, and 263 (the latter at the limit of chromosome assembly and synteny could not be confirmed). Chromosome 14 from *T. cruzi* Brazil A4 presents synteny with two scaffolds of *T. cruzi* Dm28c genome assembly: 052 and 025. Even tDNAs are useful for these determinations in chromosomes formed by the disruptive compartment, where the synteny between genomes is lost (Chr2 Brazil A4 and 027, 088 and 001 from Dm28c). Therefore, analyzing the genomic position of tDNAs contributes to generating more contiguous genome assemblies.

**Figure 7:**
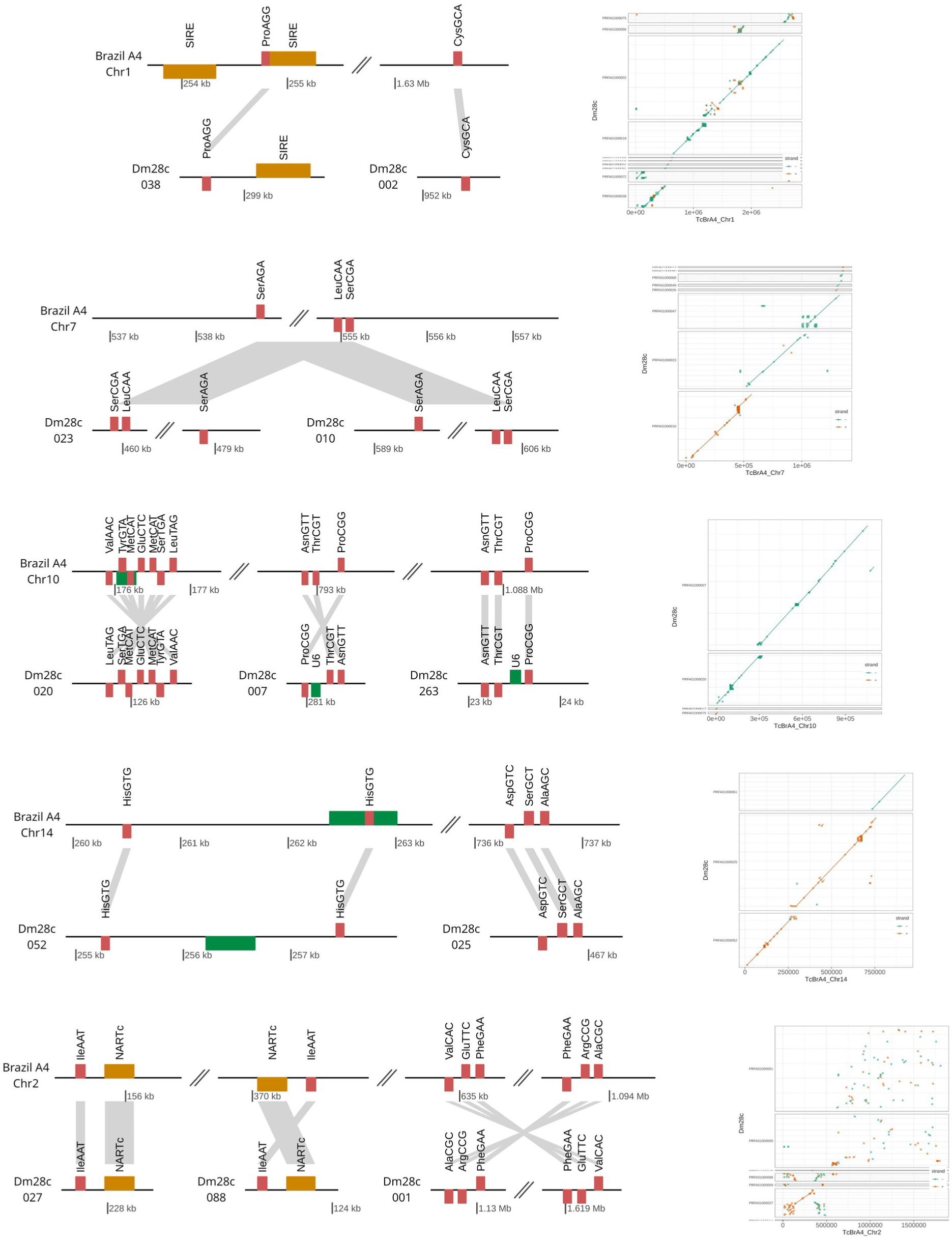
tRNA genes as markers for genome scaffolding. Genomic representation of tDNA clusters in Dm28c and Brazil A4. Grey lines represent the synteny between them. Whole chromosomal alignments

## Discussion

Most of the tRNA genes in *T. cruzi* are organized in clusters, representing a substantial difference from the rest of the eukaryotes, where tRNA genes are largely monocistronic (Maraia et al., 2017). This is also valid for the rest of the trypanosomatids since we observed synteny among the genomes in the tDNA loci. Indeed, the lack of previously identified synteny arises from misassembled genomes. This issue can now be addressed due to the improved quality of genome assemblies made possible by using long-read sequences.

Most eukaryotes exhibit tens or even hundreds of thousands of tDNA and exhibit several copy numbers within their genomes (Bermudez-Santana et al., 2010). For instance, *Saccharomyces cerevisiae* possesses 275 tRNA genes, humans have 428, and mice have 400 tRNA genes (GtRNAdb Data Release 21). In contrast, trypanosomatids genomes harbor just over a hundred copies, and the majority are single-copy genes despite being in a genomic context of remarkable plasticity, capacity for recombination, and generation of tandem repeats. Remarkably, although the *T. cruzi* genome, in particular, presents an expansion at the expense of repeated elements, multigene families, and tandem genes, the tRNA genes maintained the number of genes reduced and their organization.

tDNA loci in *T. cruzi* were described as regions where the unusual H2B histone variant (H2B.V) was preferentially located, generating unstable nucleosomes (Rosón et al., 2022). Here, we observed that tRNA genes mostly appear at the boundaries of directional gene clusters (DGCs), and several of them are present at the boundaries of CFDs. Taken together, these observations indicate that tRNA genes in *T. cruzi* are also implicated in other functions at the chromatin level.

The number of tRNA isoacceptors gives a good approximation of the abundance of each tRNA, which usually correlates with the frequency of occurrence of codons in coding DNA (Ikemura et al., 1985). There is a good correlation between codon usage and the number of tRNAs for each isoacceptor in *T. brucei* and *L. major* but not in *T. cruzi*, previously explained probably due to the use of a hybrid strain for the analysis (Padilla-Mejia et al., 2009). However, we analyzed a non-hybrid strain, and we also observed a weak correlation. This absence of correlation between these two variables also cannot be explained by the missing isoacceptors in the genome. 45 isoacceptors were identified in the Dm28c genome, and the 16 missing tRNA are the same tRNA missing in *the T. cruzi* hybrid strain, *T. brucei,* and *L. major* genome assemblies (Padilla-Mejia et al., 2009). The only difference between *T. cruzi* (hybrid and non-hybrid) and the others is the number of genes that encode each isoacceptor. In the hybrid strain, the isoacceptors are encoded mainly by two or four genes; in the non-hybrid strain, the majority are by three; meanwhile, in *T. brucei* and *L. major,* the vast majority are encoded by one gene. The correlation between codon usage and the number of individual tDNAs is usually observed in prokaryotes and not in eukaryotes, and this has been associated with additional roles of tDNAs beyond protein synthesis in eukaryotes (Iwasaki et al., 2020). The lack of correlation in *T. cruzi* raises questions about the additional roles of tDNA in this organism.

The association of tDNAs with TE had not been described in trypanosomatids genomes. In addition to TEs, the tDNA loci hold nine small nuclear ARNs (snRNAs), two 7SL RNA genes, and five sRNA76. This linear genome association of other Pol III transcribed genes was previously described in trypanosomatids and constitutes a mechanism of regulation of the transcription of these small RNAs (Nakaar et al., 1994). As in *T. brucei*, 5S rRNA genes are not clustered with tDNAs in *T. cruzi*, a phenomenon that occurs in *L. major* (Padilla-Mejia et al., 2009). Besides having similar gene composition, we found a high level of synteny in the tDNA locus among both Trypanosoma species.

A surprising finding was the high copy number of the tRNA^SeC^ genes in *T. cruzi* since they are usually present as single-copy genes in species as distant as *H. sapiens*, *D. melanogaster*, *C. elegans* and *M. musculus* (GtRNAdb Data Release 21). Furthermore, *T. brucei* and *T. cruzi* share an ancestral organization where the same two protein-coding genes flank the tRNA^SeC^ gene. The general rule found in this work that in trypanosomes, tRNA genes resist gene duplication is not fulfilled for the tRNA^SeC^ gene, which leads us to hypothesize that its extra translational functions would not be as relevant as in the rest of the tDNAs. If this is the case, its high copy number could be because the gene was swept away by a wave of gene duplication, the biological goal of which was to expand a protein-coding gene. The presence of well-positioned nucleosomes in all copies of the tDNA^SeC^ could indicate that its expression is strictly regulated at the chromatin and transcriptional levels. That is, when RNA pol II transcribes this region, the presence of nucleosomes will prevent its expression, and only the kinase will be transcribed. Future studies are necessary to confirm this hypothesis.

## Figure Legends

**Supplementary Figure 1:** Extended figure of the genomic organization of tDNAs in *T. cruzi* Dm28c including the protein-coding genes surrounding the tDNAs loci.

**Supplementary Figure 2:** Genomic organization of tDNAs in *T. cruzi,* including fragments of TE at tDNA locus.

**Supplementary Figure 3:** Secondary structures and tRNA genes length distribution.

**Supplementary Figure 4:** Multiple sequence alignment

**Supplementary Figure 5:** Multiple sequence alignment

**Supplementary Figure 6:** Comparative analysis of tRNA^Sec^ genes of trypanosomatids (*T. cruzi* and *L. major* as representatives of the genus Trypanosoma and Leishmania, respectively) with genes of other eukaryotes. **A.** The positions in the MSA that group the sequences according to genus (Supplementary Figure 4) were studied in comparison with tRNA^Sec^ from other eukaryotes. **B.** The secondary structures of *T. cruzi* and *T. brucei* were predicted by tRNAscan-SE, while the rest of the secondary structures were obtained from tRNAdb. Then, the variations observed in the tRNA^Sec^ genes identified in the trypanosomatid genomes were indicated in the secondary structures of *T. cruzi*. In blue are indicated the MSA positions that present mismatches and distinguish the sequences according to the genus.

**Supplementary Figure 7:** Genomic organization of tDNAs in *T. cruzi* TCC.

**Supplementary Figure 8:** Extended figure of the genomic organization of tDNAs in *T. cruzi* TCC, including the protein-coding genes surrounding the tDNAs loci.

**Supplementary Figure 9:** Genomic organization of tDNAs in *T. cruzi* Brazil A4. tDNA coding genes annotated using tRNA-scan SE were identified in 38 different loci in the *T. cruzi* Brazil A4 genome. 71% of the tDNAs are organized in clusters and in linear genome association with transposable elements. Protein-coding genes are indicated in green without annotation.

**Supplementary Table 1**: tDNAs identified in *T. cruzi* Dm28c genome.

**Supplementary Table 2:** Structural characteristics of tDNA identified in the *T. cruzi* Dm28c genome.

**Supplementary Table 3:** tRNA^SeC^ genes identified in trypanosomatids genomes.

**Supplementary Table 4:** tDNAs identified in *T. cruzi* TCC genome.

**Supplementary Table 5:** Characteristics of tDNAs of *T. cruzi* Dm28c and codon usage.

**Supplementary Table 6:** tDNAs identified in *T. cruzi* Brazil A4 genome.

**Supplementary Table 7:** tDNAs identified at CFD boundaries in *T. cruzi* genome.

**Supplementary Table 8:** Well-positioned nucleosomes at tDNA^SeC^ loci in *T. cruzi* genome.

**Supplementary Table 9:** Genome features identified at CFD boundaries in *T. cruzi* genome.

## Methods

tRNAScan-SE v2.0.3 (Lowe and Eddy 1997) Eukaryotic mode was used for tDNAs-prediction on *T. cruzi* Dm28c (TcruziDm28c2018 from TriTrypDB version 45), *T. cruzi* Brazil A4 (TcruziBrazilA4 from TriTrypDB version 53), *T. cruzi* TCC (TcruziTCC from TriTrypDB version 63), *T brucei* (Tbruceibrucei927 from TriTrypDB version 45) and *L. major* (LmajorFriedlin from TriTrypDB version 45) genome assemblies. Genome FASTA and GFF files were retrieved from TritrypDB (https://tritrypdb.org/tritrypdb/app) (Aslett et al., 2010).

The genome context analysis and representation were performed with genoplotR (Guy et al., 2010). tDNAs were assigned to different genomic regions (core or disruptive) using intersect function of BEDTools (Quinlan et al., 2010) v2.27.1. tDNA loci were defined as regions of the genome between two protein-coding genes containing at least one tDNA gene and a tDNA cluster as two tDNAs located in a tDNA loci.

To identify 7SL genes in the genomes, known sequences from *T. brucei* and *Trypanosoma congolense* (Michaeli et al., 1992; Chiweshe et al., 2019) were used to generate a covariance model. In the case of snRNAs, we created covariance models with sequences available in RNAcentral (https://rnacentral.org/). The sequences were aligned with T-Coffee (Notredame et al., 2000), adding the consensus secondary structure (alifold from the Vienna package). The alignments output in Stockholm format was used to build the covariance models using the CMBUILD function from the INFERNAL (Nawrocki et al., 2013). The calibrated models were then used to search the genomes using the CMSEARCH. Two hits with very good scores were obtained for 7SL, 2 for snRNA U1, 5 snRNA U2, 2 snRNA U3, 1 snRNA U4, 1 snRNA U5, and 2 snRNA U6.

Codon usage (codon relative frequencies) was calculated by using uco function from seqinr package (Charif et al., 2007) in R. To determine if there were significant differences, Spearman’s correlation coefficient was used.

To analyze the synteny, the genome assemblies were compared using MUMmer (Kurtz et al., 2004) v3.0 alignment tool. The minimum length of a single match and the minimum length of a cluster of matches were set to 100 and 500 nt, respectively in the nucmer function. After comparing whole genomes, the delta file was filtered (delta-filter -l 10000 -q -r) to get only the contigs with the best alignments. The results were visualized using ggplot2 in R.

Multiple sequence alignments (MSA) were conducted using MAFFT (Katoh et al., 2013) and visualized in Jalview (Waterhouse et al., 2009). The conserved positions in all the sequences are indicated in color.

RNA-seq, MNaseq, and Hi-C datasets were analysed as previously described (Díaz-Viraqué et al., 2023). A GFF file of CFD boundaries for the *T. cruzi* genome was obtained from the Supplementary Data of Díaz-Viraqué et al (Díaz-Viraqué et al., 2023). We compared the tRNA genes genomic coordinates with CFD boundaries +/- 5kb using the intersect function of BEDTools (Quinlan et al., 2010) v2.27.1.

All figures were designed using GIMP v2.10.32.

## Author contributions

C.R. and R. E. designed the research; F.D.V. performed the research and analyzed data, F.D.V., R. E., and C.R. wrote the manuscript.

## Acknowledgments

This work was supported by funding from GCRF (“A Global Network for Neglected Tropical Diseases” MR/P027989/1); Universidad de la República (Uruguay), CSIC Iniciacion 22320200200121UD (FDV); Agencia Nacional de Investigación e Innovación (ANII, Uruguay), Proyecto de Investigación Fundamental “Fondo Clemente Estable” FCE_3_2022_1_172653 (FDV); and FOCEM (COF 03/11). FDV received fellowships from the ANII (POS_NAC_2016_1_129916) and CAP (Universidad de la República, BFPD_2021_1#45569540). FDV, RE, and CR are National System of Researchers (SNI-ANII, UY) members. CR and RE are PEDECIBA (Programa de Desarrollo de Ciencias Básicas, Uruguay) researchers. We want to thank Laboratorio de Interacciones Hospedero-Patógeno-UBM (IPMontevideo) members for their helpful comments and many interesting discussions.

